# High school science fair: Experiences of two groups of undergraduate bioscience students

**DOI:** 10.1101/2021.01.10.426070

**Authors:** Frederick Grinnell, Simon Dalley, Joan Reisch

## Abstract

Science fairs offer potential opportunities for students to learn first-hand about the practices of science. Over the past six years we have been carrying out voluntary and anonymous surveys with regional and national groups of high school and post high school students to learn about their high school science fair experiences regarding help received, obstacles encountered, and opinions about the value and impact of science fair. Understanding what students think about science fairs will help educators make science fairs more effective learning opportunities. In this paper, we focus on the findings with two national groups of post high school students – undergraduate research fellows (SURF students) who did research at UT Southwestern Medical Center during 2014-2019 and undergraduates biology students attending the 2019 Howard Hughes Medical Institute Science Education Alliance (SEA) summer symposium. About 25% of the students who completed surveys indicated that they had participated in high school science fair, but more than half attended high schools where science fairs were unavailable. Effectively, 6 out of every 10 students participated in science fair if available. Students who could have participated in high school science fair but chose not to do so identified *not enough time* and *coming up with their project idea* as major reasons why not. About half the SURF students favored requiring non-competitive science fair regardless whether they themselves had participated in science fair. On the other hand, less than 1 in 5 thought that competitive science fair should be required. *Introduction to the scientific process* and *general learning* were mentioned most frequently as the reasons to require non-competitive science fair; these reasons were mentioned rarely in connection with competitive science fair. Unlike the national cohort of high school students we surveyed previously, who mostly did science fair in 9^th^ and 10^th^ grades, SURF students participated in science fair throughout high school and were twice as likely as high school students to have carried out science fair more than once. In conclusion, our findings suggest that participation of the undergraduate bioscience majors in high school science fairs occurs far more frequently than recognized previously. The findings emphasize further the importance of incentivizing rather than requiring science fair participation, especially in 9^th^ and 10^th^ grades, and the potential value of developing non-competitive science fairs.

## Introduction

“Science investigation and engineering design should be the central approach for teaching and learning science and engineering,” according to recommendation #1 of the 2019 National Academies Report *Science and Engineering for Grades 6-12* [1]. The idea that students should experience for themselves the eight practices of science and engineering [2] has become an underlying assumption of *Next Generation Science Standards* (NGSS) [3]. How to achieve this goal as a practical matter remains uncertain [4].

Since their origination almost 100 years ago, science fairs have come to attract a lot of public attention [5]. Some educators suggest that science fairs offer an ideal opportunity for students to experience the eight practices of science and engineering [6-10]. However, research aimed at determining if science fair participation in high school encourages science education engagement post high school suggested only a small effect. According to the national OPSCI survey of students, participation in a STEM competition was connected with only a 5% greater likelihood of STEM career interest at the end of high school [11]. These findings and an earlier study [12] concluded that the major impact of STEM competitions was to help retain students already interested in STEM rather than attract those previously uninterested. In the Microsoft Corporation STEM perceptions study, only about 5% of the students said that they became interested in STEM because of science fairs and contests [13]. In a study of students at a Queensland science and engineering university, 7% listed science fair as the reason they became interested in STEM [14]. However, high school seniors from a Texas charter school system exhibited no significant difference in STEM interest that could be attributed to participation in science fair [15].

Several years ago, we began an ongoing research program to learn what students thought about their science fair experiences, a question about which little was known. Our overall goal has been to identify strengths and weaknesses and potential improvements that might enhance science fair learning outcomes. Using anonymous, voluntary surveys, we asked regional and national groups of high school and post high school students questions regarding help received, obstacles encountered, and opinions about the value and impact of science fair. Surveying both high and post high school students as we have done provides two potentially different views of science fairs, a “real-time” view from students who have just participated in science fair but whose possible career interests in science are not established; and a retrospective view from students who may or may not have participated in science but fair, but who are pursuing bioscience-related educational interests.

In 2015 and 2016, regional high school students were invited to participate in our surveys immediately after the Dallas Regional Science and Engineering Fair. Beginning in 2017, national cohorts of high school students were contacted indirectly by incorporating our surveys into the web-based application called *Scienteer*, now adopted by Texas, Alabama, Maine, Missouri, Vermont, and Virginia for student science fair registration, parental consent, and project management. Students had the option to complete a survey after they had finished completely all science fair competitions. Unlike the regional group of high school students, of whom only 6% (4 of 64 students) were required to participate in science fairs [16], 67% (245 of 363 students) of the students in the 2017 & 2018 national cohorts indicated that they were required to participate in science fairs [17]. Regardless of their interests in science and engineering, regional and national high school students were opposed to being required to participate in science fair [17, 18]. Moreover, students required to participate in science fairs reported reduced interest in the sciences or engineering and an increased likelihood of engaging in research misconduct [17]. Required participation in science fair had not been recognized previously as an important variable regarding the impact of science fair. Our research provides empirical evidence that as best practices, student participation in high school science fair should be incentivized rather than required [19] as has been recommended by the National Science Teaching Association [20].

The post high school students that we have surveyed beginning in 2014 are on bioscience education trajectories and consist of UT Southwestern Medical Center (UTSW) medical students (mostly from Texas) doing summer research projects, biomedical science graduate students (an international group), and undergraduate research fellows doing summer research (SURF students). The SURF students are rising juniors, ∼75 of whom visit UTSW every summer from U.S. colleges and universities. About one quarter of the post-high school students who completed surveys in 2014 and 2015 had participated in high school science fair [16]. Why more post high school students did not participate in science fairs could have resulted from diverse reasons including lack of access. For instance, the National Center for Education High School Longitudinal Study of 2009 reported that only about 1/3 of the 900 schools surveyed “holds math or science fairs, workshops or competitions” [21]. To learn more about the reasons why post high school students did not participate in science fairs, we added new survey questions beginning in 2016.

Findings in this paper are based on surveys completed by SURF students over the period 2014-2019 (286 students completed surveys). To increase the potential robustness of the findings, we also surveyed a second post high school group in 2019, students who attended that years’ Howard Hughes Medical Institute (HHMI) annual Science Education Alliance (SEA) summer symposium (53 students completed surveys). The HHMI SEA program involves biology departments across ∼200 U.S. colleges and universities. Each campus can send one student to participate in the symposium. For both the SURF and HHMI student groups, we learned that ∼25% of the students who completed surveys participated in high school science fair, consistent with the preliminary findings, but we learned that more than half of the SURF students attended high schools where science fairs were unavailable. Effectively, therefore, 6 out of every 10 students participated in science fair if available. Comparison of the SURF students with the national cohort of high school students showed some important differences in student experiences. Unlike the high school students who mostly did science fair in 9^th^ and 10^th^ grades, SURF students participated in science fairs throughout high school and were twice as likely as high school students to have carried out science fair more than once. Moreover, SURF students were much more positive about requiring non-competitive science fairs than high school students although equally opposed (5:1) to requiring competitive science fairs. These results emphasize further the importance of incentivizing rather than requiring science fair participation and the potential value of developing non-competitive science fairs, especially for 9^th^ and 10^th^ grade students. Details are reported herein.

## Materials and methods

This study was approved by the UT Southwestern Medical Center IRB (#STU 072014-076). Study design entailed administering to students a voluntary and anonymous online survey using the REDCap survey and data management tool [22]. Survey recipients were UTSW SURF students over the period 2014-2019 and biology undergraduates who participated in the 2019 HHMI SEA summer symposium. SURF students are rising juniors, ∼75 of whom visit UTSW every summer from U.S. colleges and universities. HHMI SEA students are participants in the HHMI SEA program that involves biology departments across ∼200 U.S. colleges and universities. Each campus can send one student to participate in the summer symposium. For SURF students at UTSW, survey access was accomplished by providing REDCap with student UTSW email addresses. REDCap then sends the students survey invitations directly. For undergraduates attending the 2019 HHMI SEA summer symposium, the HHMI program office provided the students with a public online link to the REDCap survey. No incentives were offered for participation.

Survey content was the same as that used previously [18] except modified beginning in 2016 to include questions about (i) whether students had science fairs available at their high schools, and (ii) if they could have participated in science fair but chose not to do so, then why not. The survey can be found in supporting information (S1 Survey), and the complete survey data set can be found in supporting information (S1 Dataset).

Quantitative data were analyzed by frequency counts and percentages. Data were sorted to compare different answer selections. Significance of potential relationships between data items was assessed using relevant statistical methods, e.g., Chi-square contingency tables for independent groups. Results shown in the figures are presented two ways -- graphically to make overall trends easier to appreciate and in tables beneath the graphs to show the actual numbers. A probability value of 0.05 or smaller was accepted as statistically significant but actual p values are indicated.

Qualitative text analysis for the open-ended text questions was accomplished as described previously [18] using an approach modeled on NVivo [23, 24] based on grounded theory [25]. Approximately 80% of the students who completed surveys wrote comments about why science fairs should be optional or required. Two members of the research team (FG and SD) independently coded students’ comments, which were categorized into a matrix of shared student reasons (nodes). The independently coded matrices were revised and harmonized into 15 categories why science fair should be required or optional. Longer student comments sometimes expressed more than one reason why, in which case the comments were coded into more than one category. As a result, the number of reasons counted exceeded the total number of student comments. The complete set of student answers to the *Reason Why* question and corresponding category assignments can be found in supporting information (S2 Dataset).

Some of the results for SURF students are compared to previously reported findings with a national cohort of high school students. These high school students had completed surveys that were incorporated into the web-based application called *Scienteer*, now used by Texas, Alabama, Maine, Missouri, Vermont, and Virginia for student science fair registration, parental consent, and project management. A detailed description of the student population, results and data sets from the 2017 and 2018 national high school student surveys can be found in reference [17] including supplemental information.

We refer to the SURF and HHMI SEA students as a national group of bioscience undergraduates. It should be recognized that these students may not be truly representative of a national sample since they are participating in prestigious programs, albeit coming from U.S. colleges and universities. Similarly, we refer to the 2017-2018 high school students as a national group recognizing that they represent students who participated in science fair and signed up to do so through the Scienteer website that is used by only six states. In general, we believe that by studying and comparing the science fair experiences of these two types of student groups, high school students for a real-time view of science fair experience and post high school students for a retrospective view, with both groups representing significant geographic distribution, the findings of our studies have potentially general implications.

## Results

### SURF student participation in high school science fair

Table 1 summarizes SURF student demographics and science fair participation based on surveys conducted during 2014-2019. A total of 621 students received survey invitations. 286 (46%) returned completed surveys. More females than males completed surveys. A small percentage of students indicated that they were unfamiliar with science fair. 73 students indicated that they had participated in high school science fair and, of these, 44% indicated that they were required to do so.

**Table 1.**
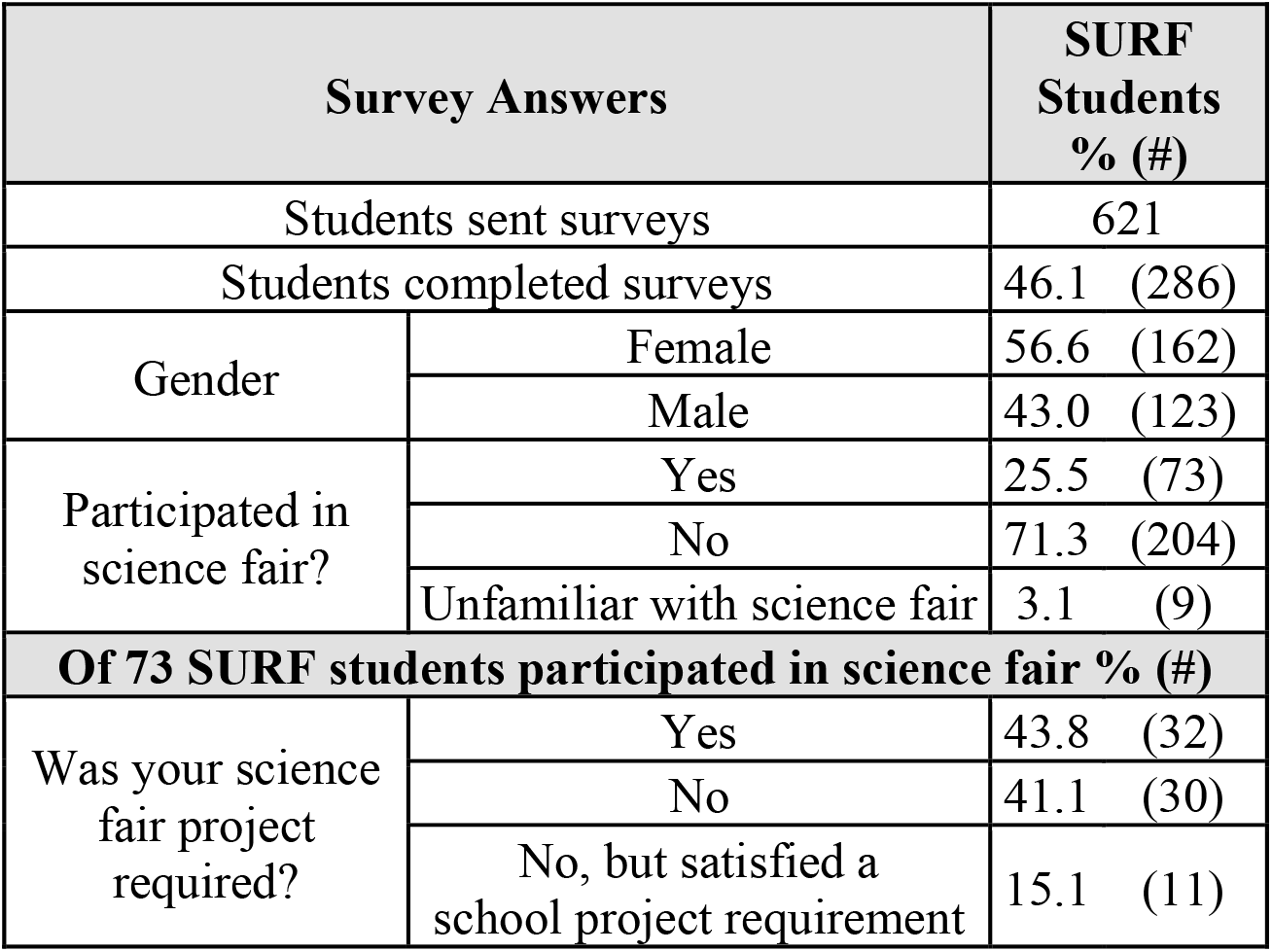
SURF student demographics and science fair requirement.

Table 2 and Figure 1 compare the SURF students with the national cohort of high school students that we surveyed during 2017 and 2018. SURF students participated in science fair to a similar extent each year of high school unlike the high school students who participated in science fair mostly in 9^th^ and 10^th^ grades [17]. Also, twice as many SURF students carried out science fair more than once compared to the high school students.

**Table 2.** SURF student science fair participation.

| Survey Answers |  | 73 SURF Students<br>% (#) | 363 HS Students<br>% (#)* | P value<br>(Chi Sq) |
| --- | --- | --- | --- | --- |
| In what grade did you carry out science fair? | 9th | 26.0 (19) | 40.5 (147) | .020 |
|  | 10th | 20.5 (15) | 32.2 (117) | .047 |
|  | 11th | 19.2 (14) | 11.3 (41) | .064 |
|  | 12th | 32.9 (24) | 11.0 (40) | <.001 |
| Carried out science fair more than once |  | 63.0 (46) | 31.1 (113) | <.001 |
| Science fair should be required if... | Non-competitive | 53.4 (39) | 25.7 (93) | <.001 |
|  | Competitive | 15.1 (11) | 20.9 (76) | .252 |

\*From [17]

**Fig.1:**
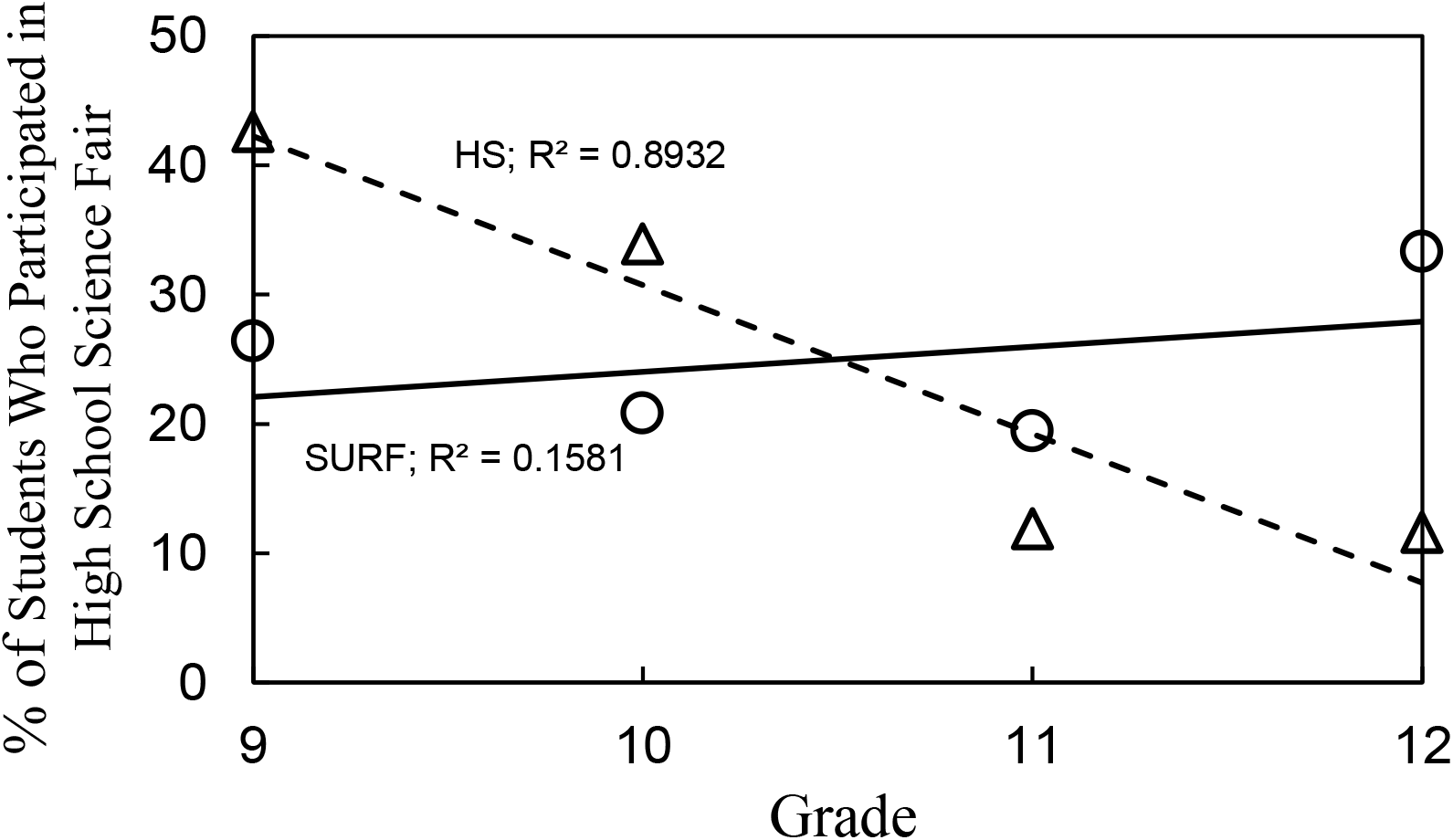
SURF vs. High school students - Grade in which participated in science fair

The findings in Table 1 and Figure 1 raised the possibility not previously considered that high school students who participated in science fair throughout all of high school might have different interests than those who participated mostly in 9^th^ and 10^th^ grades. Although we described high school student interest in science and the impact of science fair, we did not analyze grade level differences [17]. Reanalyzing the previous dataset, Figure 2 shows the grade level comparison for student interest in a career in the sciences or engineering and the impact of science fair participation on interest. Only 50% of the 147 high school students who participated in science fair in 9^th^ grade said that they were interested in a career compared to 80% of the 40 students who participated in science fair in 12^th^ grade. The percentage of high school students uninterested in a career remained the same all four years, but the percentage of unsure students dropped from 35% in 9^th^ grade to 3% in 12^th^ grade. In parallel, 55% of the high school students who participated in 9^th^ grade said that their science fair experience increased their interest in the sciences or engineering compared to 80% of the students who participated in science fair in 12^th^ grade.

**Fig. 2:**
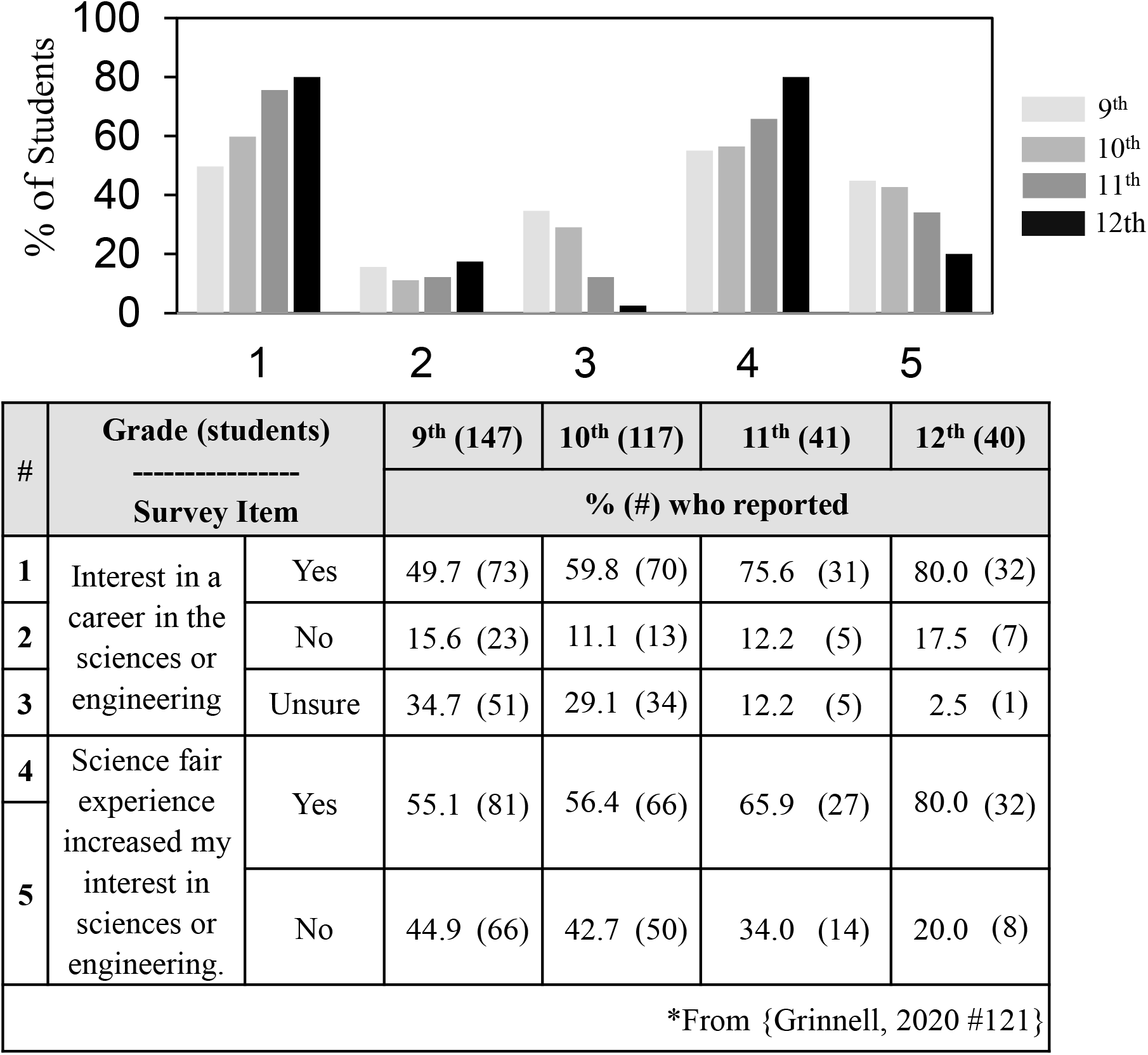
High school students - Differences in career interests and impact of science fair depending on grade in which students participated in science fair.

### SURF student reasons why science fair should be optional or required

Table 2 also shows that more than 50% of the SURF students indicated that non-competitive science fairs should be required in high school but opposed by 5:1 requiring competitive science fairs. By contrast, the national group of high school students was less inclined to favor either type of science fair requirement. This difference confirms the results previously reported in which we compared the 2015-2016 regional high school students with the 2014-2015 combined (MD/PhD/SURF) group of post high school students [18].

Figure 3 summarizes answers to the “reason why” (to require or not to require science fair) question based on comments of 280 SURF students regarding non-competitive science fair and 262 students regarding competitive science fair. Because some comments contained more than one reason, the number of reasons was greater than the number of comments. For non-competitive science fair, positive reasons (#1-#7) outnumbered negative reasons (#8-#15) (190 vs. 154); for competitive science fair, negative reasons predominated (253 vs. 68). The categories of reasons differed. The students’ positive reasons about non-competitive science fair included *introduction to the scientific process* (82 times) and *general learning* (37 times); these reasons were mentioned only 11 times total with regard to competitive science fair. *Competition incentive* was the only positive value that more students mentioned for competitive science fair compared to non-competitive science fair. Only two categories of negative reasons were similar regarding both competitive and non-competitive science fair – *negative student behaviors and consequences* and *not everyone interested in science*. Previously, we reported answers to the “reason why” question by the national group of high school students, Figure 7 in reference [17], which showed a similar overall pattern of negative reasons regarding competitive science fairs but far fewer positive reasons for requiring non-competitive science fair (131 positive vs. 314 negative).

**Fig. 3:**
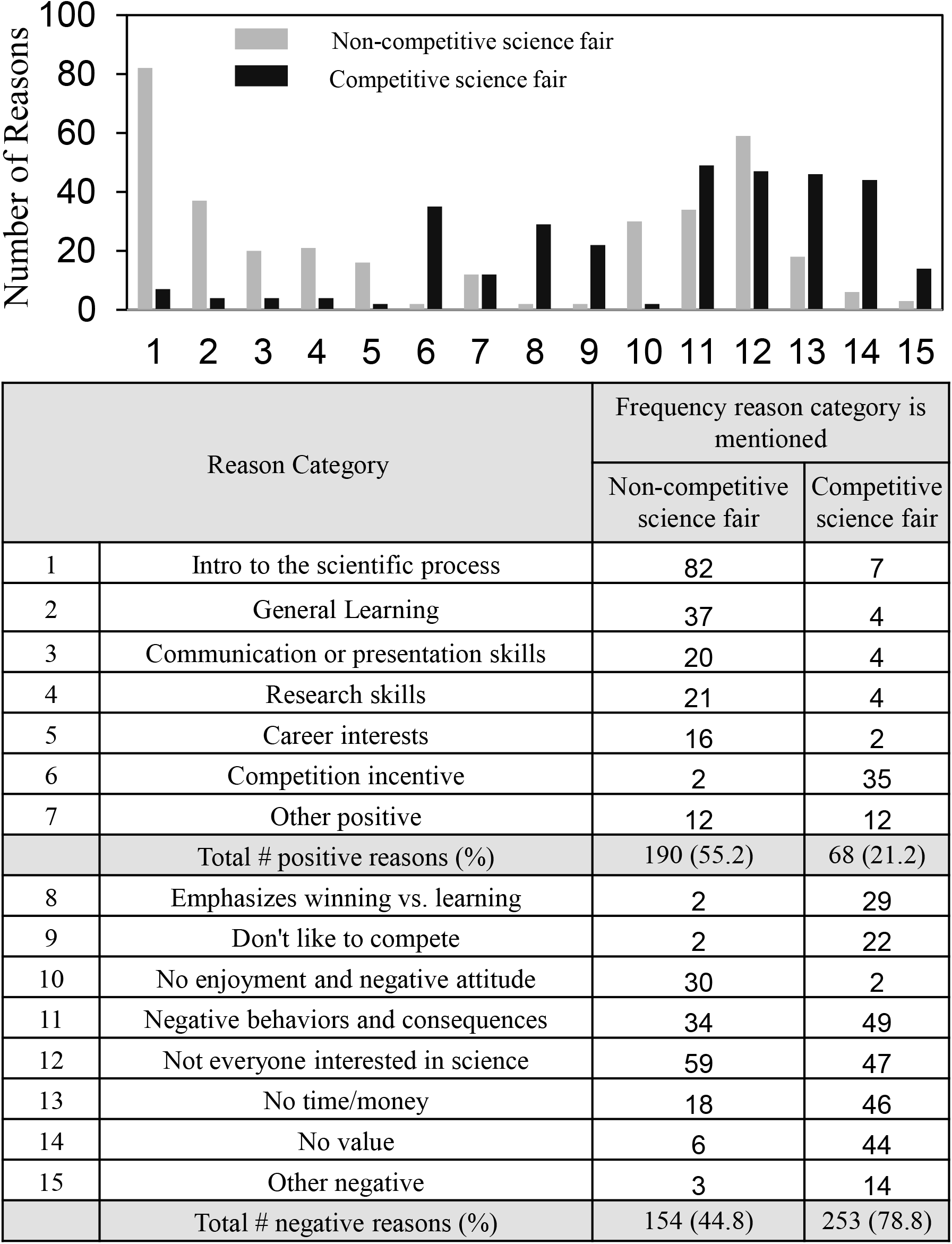
Students reasons why science fair should be optional or required

### Overall SURF student science fair experiences

Figure 4 presents a graphical summary of 2014-2019 SURF student survey answers to questions regarding sources of help, types of help received, obstacles encountered, and ways of overcoming obstacles. The corresponding supplemental figures S1-S4 show the details. Selections made by >40% of the students are labeled. These were: (A) sources of help -- parents, teachers, articles on the internet and articles from books and magazines; (B) types of help received -- developing the idea and fine tuning the report; (C) obstacles faced -- getting the idea, limited resources, limited knowledge, limited skills, and time; (D) overcoming obstacles -- picked a familiar topic, do more background research, and perseverance. None of the students indicated that they used someone else’s data (D, #12), but 4 students (5.5%) said they made up their data (D, #13) (see S4 Fig).

**Figure 4:**
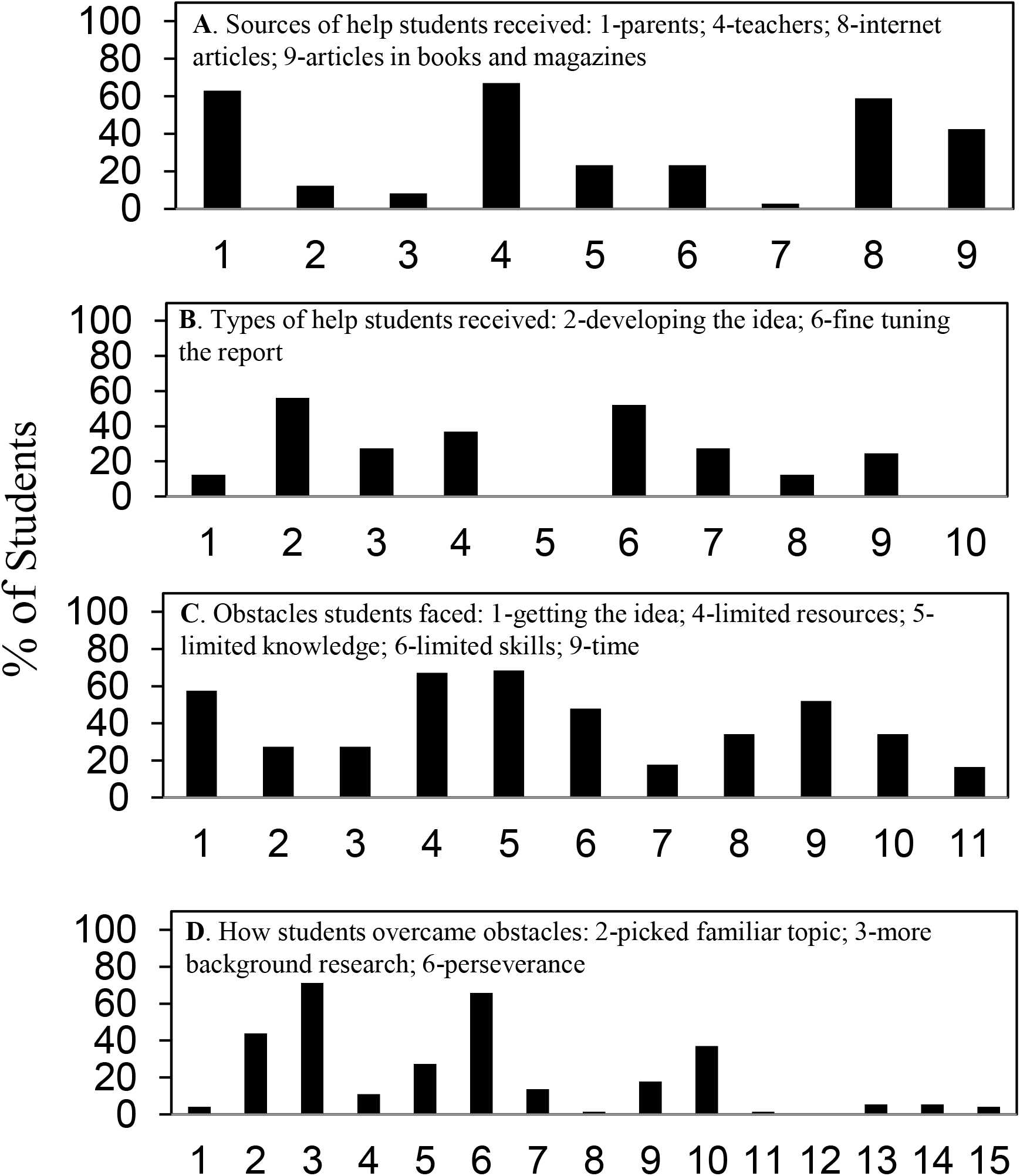
SURF summary of student experiences regarding help and obstacles

Comparison of the SURF students’ experiences with similarly graphed national high school students’ science fair experiences, Figure 2 in reference [17], showed overall similarity but some important differences. Table 3 shows that SURF students reported access to more sources of help, received more types of help, and utilized more ways to overcome obstacles. On the other hand, the SURF students reported limited resources, knowledge, and skills as obstacles twice as frequently as the high school students.

**Table 3.** Surf students vs. high school students – differences in science fair experiences.

| Survey Answers |  | 73 SURF<br>Students<br>% (#) | 363 HS<br>Students<br>% (#)* | P value<br>(Chi<br>Sq) |
| --- | --- | --- | --- | --- |
| Sources of help received | Used books or magazines | 42.5 (31) | 24.0 (87) | .001 |
| Types of help received | Development of the idea | 56.2 (41) | 26.2 (95) | < .001 |
|  | Fine tuning the report | 52.1 (38) | 32.2 (117) | .001 |
| Obstacles faced | Limited resources | 67.1 (49) | 36.1 (131) | < .001 |
|  | Limited knowledge | 68.5 (50) | 30.6 (111) | < .001 |
|  | Limited skills | 47.9 (35) | 22.0 (80) | < .001 |
| Ways to overcome obstacles | Picked familiar topic | 43.8 (32) | 29.8 (108) | .019 |
|  | More background research | 71.2 (52) | 48.5 (176) | < .001 |
|  | Perseverance | 65.8 (48) | 44.4 (161 ) | < .001 |
| *From [17] |  |  |  |  |

### SURF student science fair experiences and gender differences

Our previous studies with high school students did not show gender-dependent differences in the students’ science fair experiences [16]. However, some differences were evident for the SURF students. Table 4 shows that females experienced getting motivated as an obstacle less frequently than males, but females were more likely to report limited resources as an obstacle. Males were more likely than females to stop working on their projects as a way of overcoming obstacles, whereas females were more likely to have someone else keep them on track. Four students (about 5% of all those surveyed), all males, reported that they made up their data.

**Table 4.** Gender differences in SURF student science fair experience.

| Survey answers by students who participated in science fair |  | Males<br>33 students<br>% (#) | Females<br>40 students<br>% (#) | p Value<br>(Fisher) |
| --- | --- | --- | --- | --- |
| What obstacles did you encounter? | Getting motivated | 45.4 (15) | 12.5 (5) | .003 |
|  | Limited resources | 54.5 (18) | 77.5 (31) | .048 |
| How did you overcome obstacles? | Stopped working on the project for awhile | 21.2 (7) | 2.5 (1) | .020 |
|  | Had someone else keep me on track | 3.0 (1) | 22.5 (9) | .019 |
|  | Made up the data | 12.1 (4) | 0 (0) | .038 |
| Survey answers by students who did not participate in science fair |  | Males<br>86 students<br>% (#) | Females<br>117 students<br>% (#) | p Value<br>(Fisher) |
| What obstacles do you think students who do science fair will encounter? | Getting motivated | 62.8 (54) | 60.7 (71) | .773 |
|  | Limited resources | 76.7 (66) | 80.3 (94) | .603 |
| How do you think students who do science fair will overcome obstacles? | Stop working on the project for awhile | 16.3 (14) | 14.5 (17) | .843 |
|  | Have someone else keep on track | 44.2 (38) | 49.6 (58) | .479 |
|  | Make up the data | 23.3 (20) | 19.7 (23) | .603 |

We also asked students who did not participate in science fair to answer hypothetically the questions about obstacles encountered and overcome. Table 4 (bottom section) shows that students who did not participate in science fair anticipated more obstacles and more ways to overcome obstacles but no apparent gender differences in their responses. About 20% of the students who did not participate in science fair anticipated that students would make up their data, which was consistent with our earlier findings [16].

### SURF students’ opportunity to participate in high school science fair and reasons not to participate

The effective rate of SURF student participation in high school science fair at 25% (Table 1) could have been higher since it did not take into account whether science fairs were available to the students. For instance, the High School Longitudinal Study of 2009 reported that only about 1/3 of the 900 schools surveyed “holds math or science fairs, workshops or competitions” [21]. Table 5 shows that of the 180 students who completed our surveys during 2016-2019, which included the new questions about science fair availability, more than 50% attended high schools where science fair was <u>not</u> available. Overall therefore, the effective rate of science fair participation by the SURF students was almost 60%. Students who could have participated in science fair but did not do so most frequently chose *not enough time* and *did not have a good idea for a project* from the list of the six choices shown in Table 5 as reasons not to participate. However, students who chose not to participate in science fair were just as likely as those who participated to favor requiring non-competitive science fair.

**Table 5.** Availability of high school science fair to SURF students and reasons not to participate.

| Survey Answers |  | Students<br>% (#) |  |
| --- | --- | --- | --- |
| Students sent surveys |  | 404 |  |
| Students completed surveys |  | 44.6 (180) |  |
| Did you participate in science fair? | Science fair not available at my school | 55.6 | (100) |
|  | Could have participated but chose not to do so | 18.3 | (33) |
|  | Yes | 25.0 | (45) |
| Reasons selected by 33 students who chose not to participate in science fair even though it was available |  |  |  |
| Not enough time |  | 57.6 | (19) |
| Did not have a good idea for a project |  | 51.5 | (17) |
| Not interested |  | 33.3 | (11) |
| Did not want to compete |  | 24.2 | (8) |
| Not enough help available |  | 21.2 | (7) |
| Too expensive |  | 3.0 | (1) |
| Student attitudes about requiring science fair depending on opportunity to participate |  |  |  |
| Students say science fair should be required if: | Participated in SF<br>45 students<br>% (#) | Chose not to participate in SF<br>33 students<br>% (#) | Participating in SF not an option<br>100 students<br>% (#) |
| Non-competitive | 51.1 (23) | 48.5 (16) | 37.3 (38) |
| Competitive | 17.8 (8) | 12.1 (4) | 6.9 (7) |

### HHMI SEA students’ opportunity to participate in high school science fair

UT Southwestern SURF students represent a national cohort of undergraduates coming from schools all over the United States. Nevertheless, to test further the generality of our findings regarding student participation in science fairs, we surveyed another national student group, biology undergraduates attending the 2019 HHMI SEA summer symposium. The HHMI SEA program involves about 200 biology departments in U.S. colleges and universities. Each department can send one student representative to the symposium. Table 6 shows that 23% of the HHMI SEA students had participated in high school science fair; of these, 80% said that they were required to participate. However, the majority of the high schools attended by the HHMI SEA students did not offer an opportunity to participate in science fair. Therefore, overall 75% (12 of 16) of the HHMI SEA students who could have participated in science fair did so.

**Table 6.** HHMI SEA student participation in high school science fair.

| Survey Answers |  | HHMI SEA<br>% (#) |
| --- | --- | --- |
| Students sent surveys |  | 150 |
| Students completed surveys |  | 35.3 (53) |
| Did you<br>participate in<br>science fair? | Science fair not available at my school | 64.2 (34) |
|  | Could have participated but chose not to do so | 7.5 (4) |
|  | Yes | 22.6 (12) |
|  | Unfamiliar with science fair | 5.7 (3) |
| Was science<br>fair required if<br>available? | Yes | 83.3 (10) |
|  | No | 16.7 (2) |

## Discussion

Over the past six years we have been carrying out voluntary and anonymous surveys with regional and national groups of high school and post high school students to learn about their high school science fair experiences regarding help received, obstacles encountered, and opinions about the value and impact of science fair. Underlying our research is the expectation that understanding what students think about science fairs will help educators make science fairs more effective learning opportunities. Surveying both high and post high school students, as we have done, potentially offers two different views of science fairs that we can compare. On one hand, we learn a “real-time” view from students who have just participated in science fair but whose possible career interests in science are not established. On the other, we learn a retrospective view from students who may or may not have participated in science fairs, but who are pursuing STEM-related educational trajectories, in our case in the biosciences.

In this paper, we focus on findings based on 286 SURF students who completed surveys between 2014-2019 and 53 HHMI SEA students who completed surveys in 2019. SURF students are rising juniors, ∼75 of whom visit UTSW every summer from U.S. colleges and universities. HHMI SEA students are participants in the HHMI SEA program that involves biology departments across ∼200 U.S. colleges and universities. Each campus can send one student to participate in the summer symposium. Overall, 25.5% of the students and 22.6% of the HHMI SEA students indicated that they had participated in science fairs. By contrast, only about 5% of the 23,500 high school students surveyed in the High School Longitudinal Study of 2009 participated in a science competition [21]. Similarly, only about 5% of the almost 16,000 undergraduate students surveyed in the OPSCI college outreach survey of students in introductory freshman (mostly English) classes reported participating in high school science fair [11]. Therefore, the SURF and HHMI SEA student participation rate in high school science fair was five times higher than most undergraduate students. A related finding of the High School Longitudinal Study of 2009 was that only ∼1/3 of the 900 schools surveyed reported “holds math or science fairs, workshops or competitions” [21]. Only about 40% of the SURF and HHMI SEA students reported attending high schools where high school science fair was available. Overall, therefore, about 6 out of every 10 SURF and HHMI students who had high school science fair available participated in science fair.

Although we refer to the SURF and HHMI SEA students as national groups, it should be recognized that these students may not be truly representative of a broader national sample. Indeed, we have the opportunity to survey them because they are participating in prestigious bioscience programs. Also, since all of the students that we surveyed were on bioscience education trajectories, our findings cannot be generalized to undergraduate students with other STEM disciplinary interests. Some high school students participate in other kinds of competitions such as science Olympiads [26] and “reverse” science fairs in which high school students learn about college or graduate student research projects first and later present their own projects to the older students including the possibility of having the older students serve as mentors [27, 28]. Our findings do not provide any insights into the frequency of student participation or student experiences in these other types of competitions.

The SURF students who could have participated in science fair but chose not to do so indicated *not enough time* and *no good idea for a project* as the most frequent reasons why they did not participate. For regional and national high school student groups surveyed previously, *time pressure* and *coming up with the main idea* were the most frequently cited obstacles experienced by science fair students [16, 17]. Consequently, finding ways to reduce the time commitment and facilitate coming up with the idea for a project potentially will improve the science fair participation experience.

44% of the SURF students and 83% of the HHMI SEA students reported that their participation in science fair was required. For the national cohort of high school students surveyed previously, about 70% reported that science fair was required [17]. Since the survey does not provide students with ancillary information regarding what it means for science fair to be required, their answers reflect how they felt about their participation. We cannot tell if they understand *required to do science fair* differently from what their schools intended, e.g., required to participate in science fair to get into an advanced class or to increase one’s grade. In some cases, we know that science fair was required for advanced science classes based on comments by the high school students, e.g., “It made a lot of people choose not to take honors science, and it was a lot more work than thought out to be” [17]. Consequently, *required to do science fair* might mean different things at different stages of a student’s high school science education. In any case, as indicated by the student’s comment above and based on our previous findings, requiring science fair can become a disincentive to pursuing advanced science education [17].

Regardless whether the students were required to participate in science fair, chose not to participate, or did not have science fair available as an option, they uniformly opposed a competitive science fair requirement. On the other hand, the students were much more positive about requiring non-competitive science fair, which was favored by about half. Their positive “reasons why” to require non-competitive science outnumbered negative reasons 190 to 154. They mentioned *introduction to the scientific process* and *general learning* as positive reasons almost 120 times (35% of the total comments) regarding non-competitive science fair but only 11 times (less than 1% of the total comments) about competitive science fair. Indeed, the only positive reason mentioned frequently for competitive science fair (35 times) was *competition incentive*. In short, non-competitive science fair was viewed as emphasizing learning; competitive science fair as emphasizing winning. We suggest that the SURF students thinking back on their science fair experience were much more positive in retrospect about the potential value of science fair compared to the national group of high school students who had just participated in science fair (Figure 7) [17] since high school students’ negative reasons about requiring non-competitive science outnumbered positive reasons by 314 to 131.

The overall pattern of science fair experiences of the SURF students was mostly similar to the national cohort of high school students studied previously [17], but SURF students indicated more sources and types of help received and ways to overcome obstacles. Interestingly, the SURF students reported twice as often as the high school students *limited resources, limited knowledge*, and *limited skills* as obstacles encountered. One possible explanation is that given their interests in science, the SURF students attempted more ambitious science fair projects in high school. Another possibility is that the SURF students looking back from a more advanced time in their science education trajectories recognized their earlier limitations.

Another major difference between the SURF students and high school students concerned the timing and frequency of participating in science fair. Unlike the national cohort of high school students we surveyed previously, who mostly did science fair in 9^th^ or 10^th^ grade [17], SURF students participated in science fair to a similar extent throughout high school and were twice as likely as high school students to have carried out science fair more than once. Upon re-evaluating the national high school student cohort [17] according to grade level-dependent differences in students’ interests in science or engineering, we observed that students doing science fair could be divided overall into two groups. One group of students, those interested in careers in science or engineering, accounted for about half the students doing science fair in 9^th^ grade and 80% of the students doing science fair in 12^th^ grade. The other group, those unsure about careers in science or engineering, accounted for 35% of the students doing science fair in 9^th^ grade and 3% of those doing science fair in 12^th^ grade. Students not interested in careers remained constant at around 15%.

The difference in timing and frequency of science fair participation by students interested or not interested in science and engineering careers is consistent with the observation that a major impact of STEM competitions appears to be to help retain students already interested in STEM rather than attract those students previously disinterested [11, 12]. Given that increasing student interest in science represents one of the most important potential positive outcomes of science fair, in light of the high school students’ and SURF students’ experiences and views, we suggest that science fair organizers and schools should consider offering different experiences for science fair in 9^th^ and 10^th^ compared to 11^th^ and 12^th^ grades. For instance, in 9^th^ and 10^th^ grades, science fair could be non-competitive or have a non-competitive option. If competitive for all students throughout high school, then science fairs could be incentivized in 9^th^ and 10^th^ grades rather than required.

SURF students reported several subtle but potentially important gender differences. Females experienced *getting motivated* as an obstacle less frequently than males even though they were more likely than males to report experiencing *limited resources* as an obstacle. Males were more likely than females to *stop working on their projects* as a way of overcoming obstacles, whereas females were more likely to *have someone else keep them on track*. These differences suggest that participating in science fair was overall a more successful experience for the females. Consistent with this possibility, all of the SURF students who reported *making up their data* were males. Previous studies did not observe overall gender differences in high school science fair participation or outcomes [29-31]. Our finding that SURF females appeared to have a more successful science fair experience than males should be considered preliminary but worth further investigation. Biomedical fields do not have the same recruitment or diversity issues compared to some other STEM fields, especially physics, engineering, and computer science [32], so it would be interesting to learn if any gender differences in science fair were experienced by undergraduates in the other fields.

Finally, students who did not participate in science fair anticipated a higher level of obstacles compared to those who did participate but no gender differences in their responses. About 20% of the students anticipated that to overcome obstacles, high school students would resort to *making up their data* consistent with our findings several years ago with a smaller number of post high school students [16]. Promoting research integrity has become an important goal of the scientific community in recent years [33]. Perhaps education about research integrity should be incorporated into high school programs that aim to teach students about diverse topics in bioethics [34, 35] and especially in conjunction with science fair.

## Acknowledgments

We are grateful to Dr. Viknesh Sivanathan and Dr. David Asai from the Howard Hughes Medical Institute Science Education program for their help to survey the 2019 HHMI SEA summer symposium participants. Dr. Elise Christopher from the National Center for Education Statistics advised us about the *High School Longitudinal Study of 2009*. Use of REDCap survey and data management tool was facilitated by the UTSW Department of Population and Data Sciences and Clinical and Translational Science Training Program.

## Supporting information

**S1 Data set**. Excel dataset showing all of the survey questions and answers.

**S2 Data set**. Excel dataset showing the complete set of reason category assignments.

**S1 Survey**. Survey questions.

**S1 Fig. Frequency of student answers to the question “Who helped you with your science fair project?”**

**S2 Fig. Frequency of student answers to the question “What kind of help did you receive doing science fair?”**

**S3 Fig. Frequency of student answers to the question “In your science fair project, what obstacles did you face?”**

**S4 Fig. Frequency of student answers to the question “In your science fair project, how did you overcome obstacles?”**

**S1_fig.**
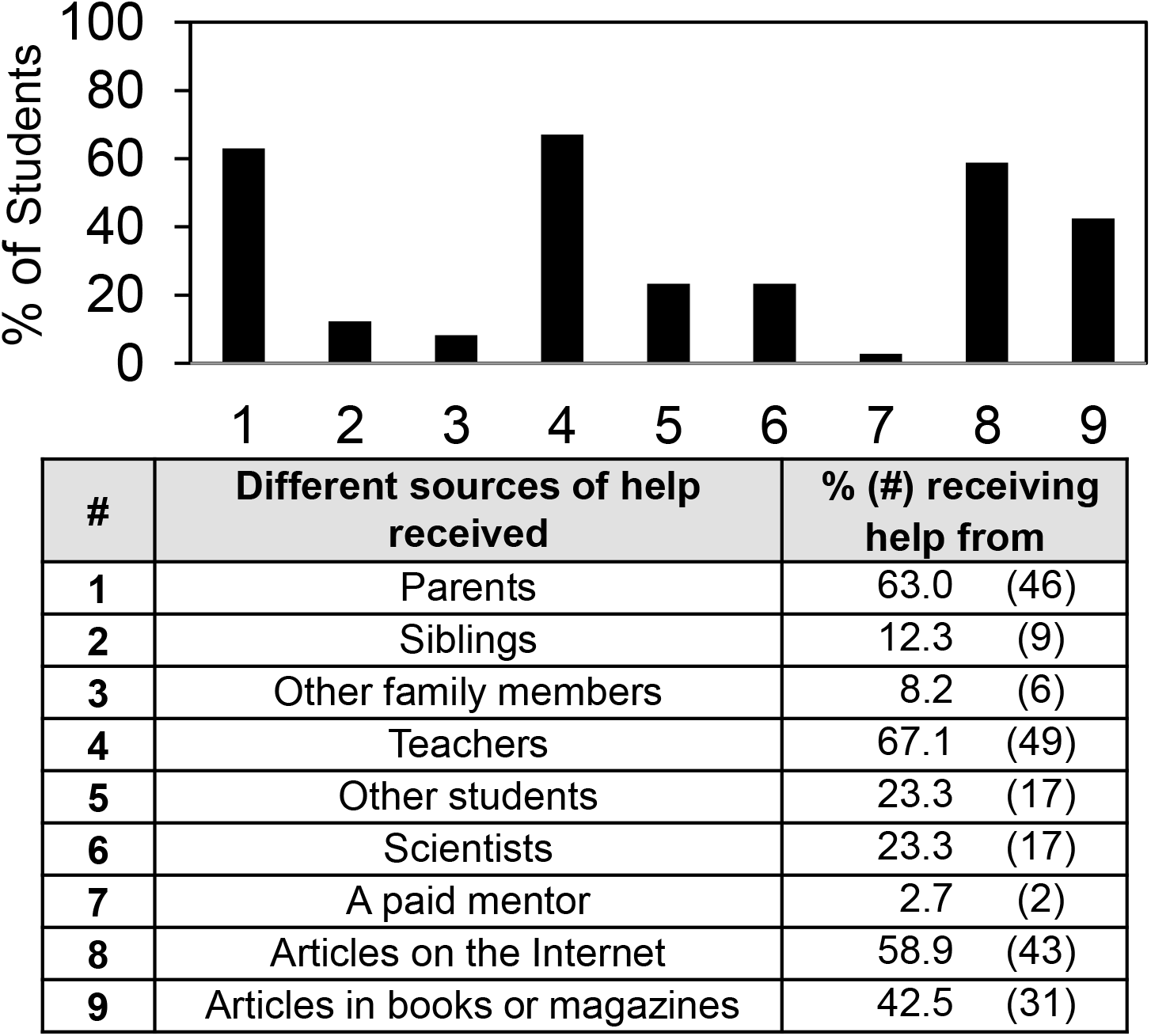
Sources of help students received in science fair.

**S2_fig.**
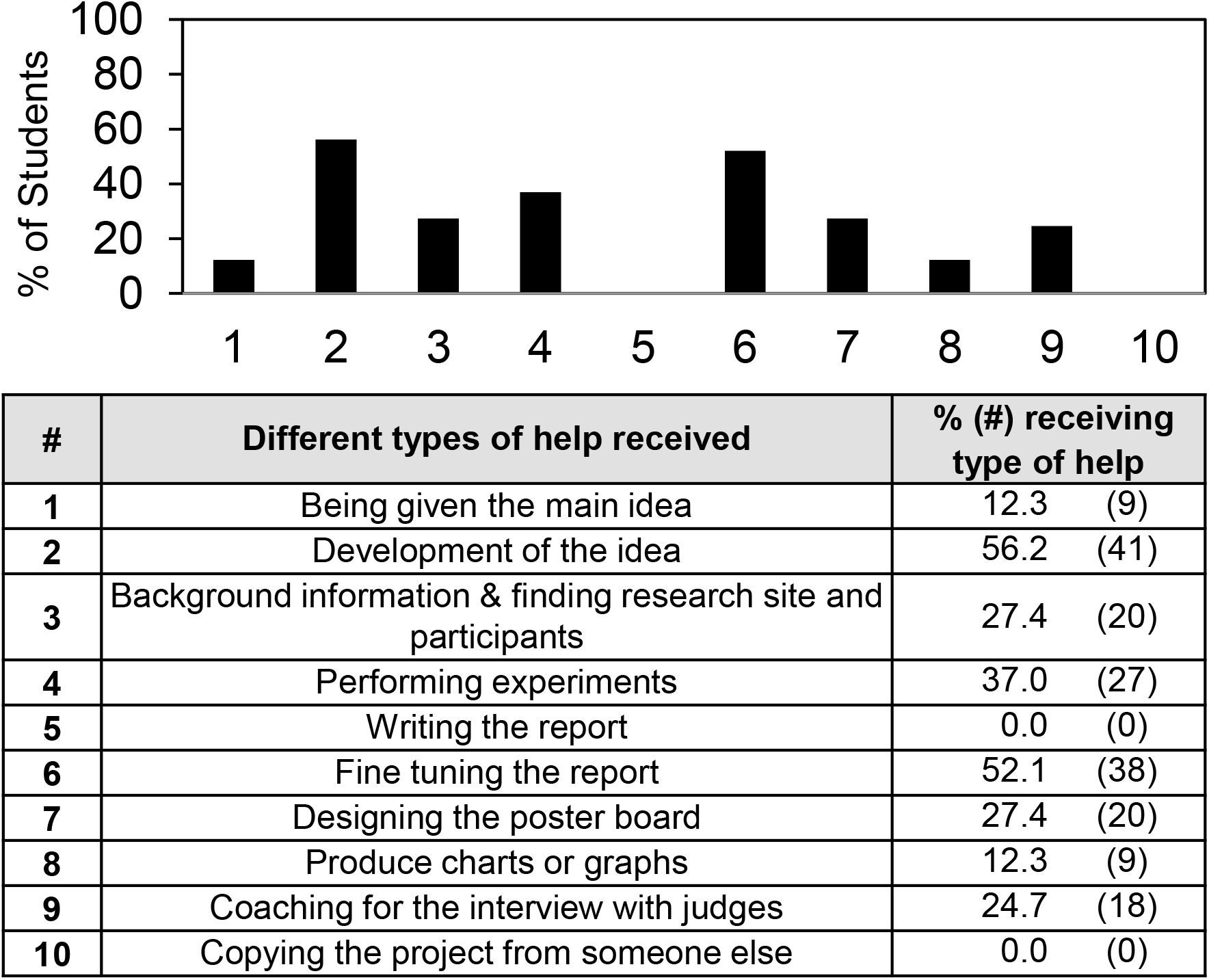
Types of help students received in science fair.

**S3_fig:**
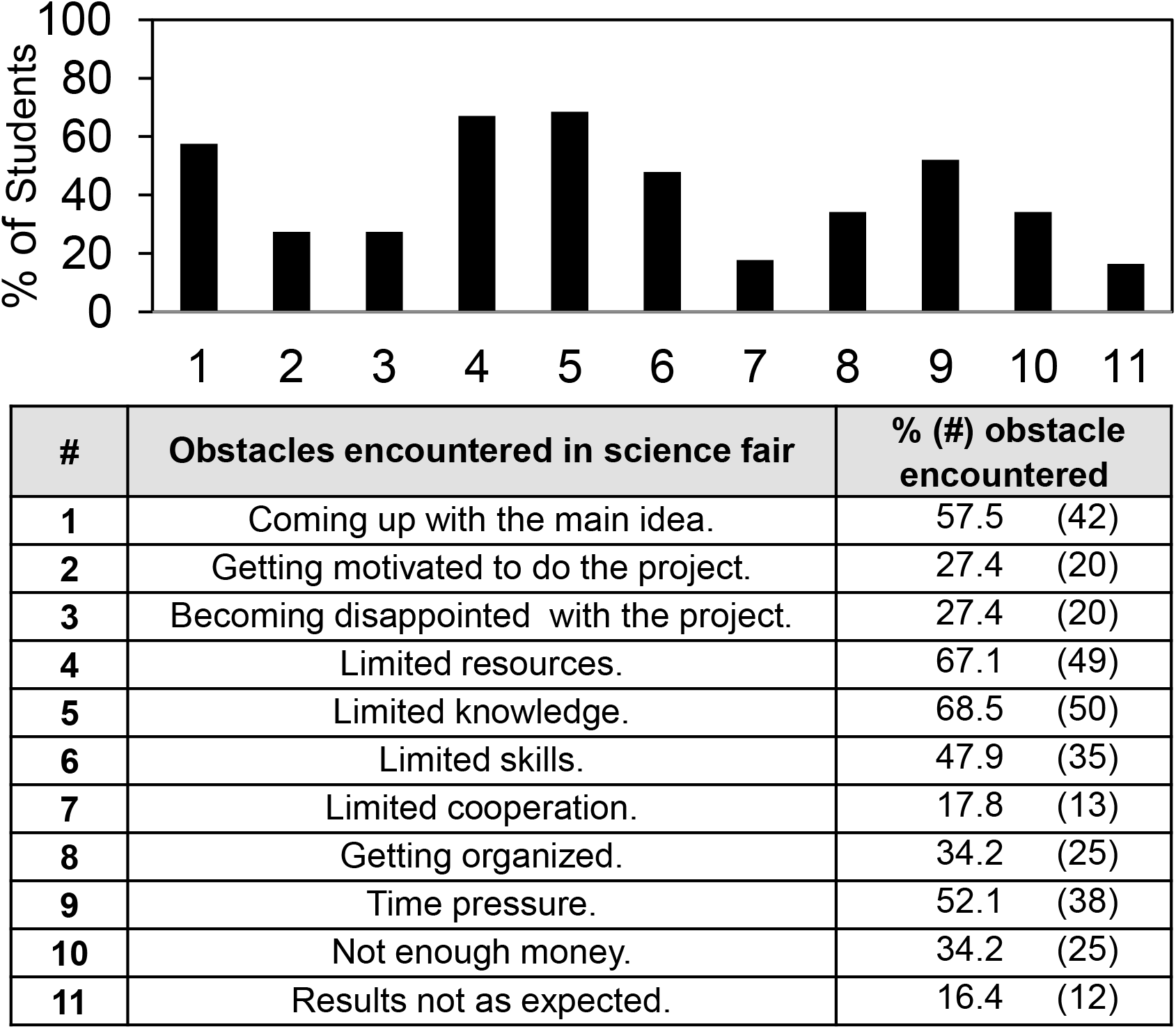
Obstacles students encountered in science fair.

**S4_fig:**
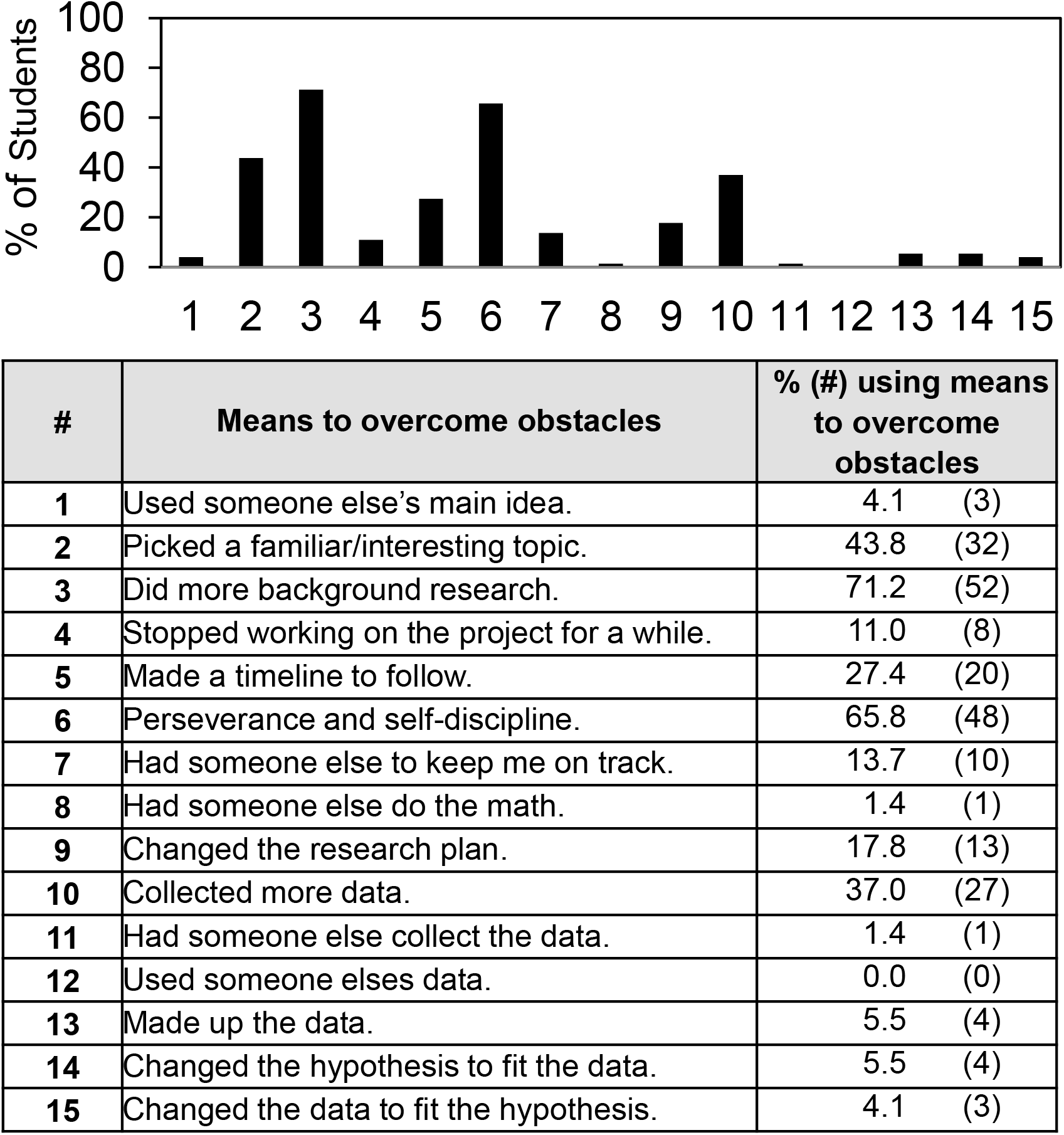
How students overcame obstacles in science fair.

